## Supplementary File 2 for "A robust method for measuring aminoacylation through tRNA-Seq"

**tRNA-charge sequencing SOP**

**Oligos, reagents and buffers**

**Oligos:**

| **Name** | **Sequence** | **IDT spec.** |
| --- | --- | --- |
| E.coli tRNA-Lys-UUU-CCA | rGrGrGrUrCrGrUrUrArGrCrUrCrArGrUrUrGrGrUrArGrArGrCrArGrUrUrGrArCrUrUrUrUrArArUrCrArArUrUrGrGrUrCrGrCrArGrGrUrUrCrGrArArUrCrCrUrGrCrArCrGrArCrCrCrArCrCrA | Ultramer RNA Oligo, 4 nmol |
| E.coli tRNA-Thr-CGT-CCAA | rGrCrCrGrArUrArUrArGrCrUrCrArGrUrUrGrGrUrArGrArGrCrArGrCrGrCrArUrUrCrGrUrArArUrGrCrGrArArGrGrUrCrGrUrArGrGrUrUrCrGrArCrUrCrCrUrArUrUrArUrCrGrGrCrArCrCrArA | Ultramer RNA Oligo, 4 nmol |
| Splint CCA | GCATGGCAGCCTGGN/3SpC3/ | DNA oligo, 250 nmole, PAGE purification |
| Splint CC | GCATGGCAGCCGGN/3SpC3/ | DNA oligo, 250 nmole, PAGE purification |
| l1Sp | /5Phos/GGCTGCCATGCGACTAAGATCGGAAGAGCACACGTCTGAA/3ddC/ | DNA oligo, 100 nmole, PAGE purification |
| l2Sp | /5Phos/GGCTGCCATGCTGTCACGAGATCGGAAGAGCACACGTCTGAA/3ddC/ | DNA oligo, 100 nmole, PAGE purification |
| l3Sp | /5Phos/GGCTGCCATGCTGCGAAGATCGGAAGAGCACACGTCTGAA/3ddC/ | DNA oligo, 100 nmole, PAGE purification |
| l4Sp | /5Phos/GGCTGCCATGCAAGCTGAGATCGGAAGAGCACACGTCTGAA/3ddC/ | DNA oligo, 100 nmole, PAGE purification |
| l5Sp | /5Phos/GGCTGCCATGCAACGCATCAGATCGGAAGAGCACACGTCTGAA/3ddC/ | DNA oligo, 100 nmole, PAGE purification |
| l6Sp | /5Phos/GGCTGCCATGCTACAGAGATCGGAAGAGCACACGTCTGAA/3ddC/ | DNA oligo, 100 nmole, PAGE purification |
| l7Sp | /5Phos/GGCTGCCATGCACATGAAGATCGGAAGAGCACACGTCTGAA/3ddC/ | DNA oligo, 100 nmole, PAGE purification |
| l8Sp | /5Phos/GGCTGCCATGCTGCGATAAGATCGGAAGAGCACACGTCTGAA/3ddC/ | DNA oligo, 100 nmole, PAGE purification |
| l9Sp | /5Phos/GGCTGCCATGCAACGTACAAGATCGGAAGAGCACACGTCTGAA/3ddC/ | DNA oligo, 100 nmole, PAGE purification |
| RT oligo | /5Phos/RNNNNNNNNNAGATCGGAAGAGCGTCGTGTAGGGAAAGAG/iSp18/GTGACTGGAGTTCAGACGTGTGCTC | PAGE Ultramer, 1 nmole |
| P7 oligo (D701) | CAAGCAGAAGACGGCATACGAGATCGAGTAATGTGACTGGAGTTCAGACGTGT*G | Regular DNA oligo, 100 nmole, standard desalting |
| P7 oligo (D702) | CAAGCAGAAGACGGCATACGAGATTCTCCGGAGTGACTGGAGTTCAGACGTGT*G | Regular DNA oligo, 100 nmole, standard desalting |
| P7 oligo (D703) | CAAGCAGAAGACGGCATACGAGATAATGAGCGGTGACTGGAGTTCAGACGTGT*G | Regular DNA oligo, 100 nmole, standard desalting |
| P7 oligo (D704) | CAAGCAGAAGACGGCATACGAGATGGAATCTCGTGACTGGAGTTCAGACGTGT*G | Regular DNA oligo, 100 nmole, standard desalting |
| P7 oligo (D705) | CAAGCAGAAGACGGCATACGAGATTTCTGAATGTGACTGGAGTTCAGACGTGT*G | Regular DNA oligo, 100 nmole, standard desalting |
| P5 oligo (D501) | AATGATACGGCGACCACCGAGATCTACACTATAGCCTACACTCTTTCCCTACACGACGCT*C | Regular DNA oligo, 100 nmole, standard desalting |
| P5 oligo (D502) | AATGATACGGCGACCACCGAGATCTACACATAGAGGCACACTCTTTCCCTACACGACGCT*C | Regular DNA oligo, 100 nmole, standard desalting |
| P5 oligo (D503) | AATGATACGGCGACCACCGAGATCTACACCCTATCCTACACTCTTTCCCTACACGACGCT*C | Regular DNA oligo, 100 nmole, standard desalting |
| P5 oligo (D504) | AATGATACGGCGACCACCGAGATCTACACGGCTCTGAACACTCTTTCCCTACACGACGCT*C | Regular DNA oligo, 100 nmole, standard desalting |
| P5 oligo (D505) | AATGATACGGCGACCACCGAGATCTACACAGGCGAAGACACTCTTTCCCTACACGACGCT*C | Regular DNA oligo, 100 nmole, standard desalting |

**Reagents:**

| **Name** | **Vendor** | **Cat#** |
| --- | --- | --- |
| TGIRT polymerase | Ingex | TGIRT50 |
| RT buffer | Thermo | 18057018 |
| KAPA HiFi DNA Polymerase | Roche | 07958854001 |
| SDS | Thermo | AM9820 |
| NaIO4 | Sigma | 311448-5G |
| TBE-Urea Gels | Thermo | EC68752BOX |
| SYBR Gold | Thermo | S11494 |
| T4 RNA ligase 2 | NEB | M0239L |
| Disposable pestles | Fisher | 12-141-364 |
| Trizol | Thermo | 15596026 |
| Lysine | Sigma | L8662-100G |
| Superase In | Thermo | AM2694 |
| rSAP | NEB | M0371L |
| ssRNA Ladder | NEB | N0364S |
| Spin filter | Sigma | CLS8161-100EA |
| CircLigase | Lucigen | CL4111K |
| Betaine | Sigma | B0300-1VL |
| DNA cleanup | Zymo | D4013 |
| Oligo purification | Zymo | D4060 |
| 10 bp DNA ladder | Thermo | 10597012 |
| Novex DNA sample buffer | Thermo | LC6678 |
| Novex TBE gel 4-12% | Thermo | EC62352BOX |
| Ethylene glycol | Sigma | 324558-100ML |
| RNaseAlert-1 Kit | IDT | 11-02-01-02 |
| Nuclease free water | IDT | 11-05-01-04 |
| RNaseZap | Thermo | AM9782 |
| NEBuffer 2 | NEB | B7002S |
| dNTPs | NEB | N0446S |

**Buffers and solvent:**

| **Buffer** | **Volume** |
| --- | --- |
| Isopropanol (IPA) | 1 L |
| 80% IPA; 50 mM sodium acetate, pH=4.5 | 100 mL |
| 100 mM sodium acetate, pH=4.5 | 10 mL |
| 200 mM NaIO4 (prepare fresh) | 1 mL |
| 50% (v/v) ethylene glycol | 10 mL |
| 1 M Lysine pH=8 (store at -20C) | 10 mL |
| TBE-Urea Sample Buffer (2x) | 10 mL |
| Gel elution buffer (0.05% SDS; 300 mM sodium acetate, pH=4.5) | 100 mL |
| 25 mM dNTP | 100 μL |
| 5 M NaOH | 1 mL |

**Preparations before RNA processing**

1. Add ethanol to wash buffers in RNA/DNA isolation kits.
2. Prepare concentrated sodium acetate solution at pH=4.5. This is best done by neutralizing a concentrated acetic acid solution with sodium hydroxide while mixing on ice. Neutralize ~45% of the acid with NaOH pellets and cool to room temperature, then adjust to final pH with dilute NaOH. Measure final volume and calculate sodium acetate concentration.
3. Sodium periodate (also referred to as sodium metaperiodate, or simply NaIO4) solution must be prepared fresh at the time of its use e.g. prepared in the morning, put on ice and used later in the afternoon. It is most convenient to prepare several 1.5 mL tubes beforehand with weighed out dry NaIO4 ready to be solubilized in 500-1000 μL water to generate a 200 mM solution (42.8 mg/mL). Keep these tubes in the freezer and take one out on the day of use. Solubilize in nuclease free water at room temperature and keep on ice until use. The same approach is advised for 100 mM DTT.
4. Prepare 10 mL of 1 M Lysine at pH=8. Adjust pH with HCl and/or NaOH. Filter through 0.45 μm filter and aliquot into 1.5 mL tubes. Lysine solutions can host bacterial growth; therefore, freeze these tubes and thaw on the day of use.
5. Prepare TBE-Urea Sample Buffer containing: 8 M Urea, 30 mM sodium acetate, 2 mM EDTA and 0.02% (w/v) bromophenol blue and xylene cyanol. Adjust pH between 4.7 and 5.
6. Prepare annealed adapter:splint partial duplex. Reconstitute the adapters and the two splint oligos in water at 100 μM. Make 20 μM adapter:splint solution using equimolar CCA and CC splint in 1x NEBuffer 2. Heat to 94°C for 2 min followed by cooling 0.3°C/s to 4°C. Store in the freezer and do not use a heated incubator to thaw.
7. Optional: gel purification of the E.coli tRNA-Lys-UUU CCA control oligo. This removes truncated oligos, resulting in higher quality. It can be done using Novex 10% polyacrylamide/7M urea/1xTBE gel, by loading 4 μL at 0.5 μg/ μL per well. Run 30 min at 100V followed by 300V until the first dye front reaches the bottom of the gel, then stain, isolate and elute from gel.
8. Optional, but highly recommended: Prepare E.coli tRNA spike-in; this controls for completion of the oxidation, cleavage and ligation reactions. It is composed of 5 ng/μL of each E.coli tRNA-Lys-UUU CCA, E.coli tRNA-Thr-CGT CCA**-Phos** and E.coli tRNA-Thr-CGT CCA (i.e. 15 ng/μL total RNA). The latter two are generated from the E.coli tRNA-Thr-CGT CCAA oligo by periodate oxidation and lysine cleavage:
   a. Reconstitute E.coli tRNA-Thr-CGT CCAA oligo in water to 100 μM.
   b. Transfer 30 μL of the oligo to 10 μL 100 mM sodium acetate pH=4.5.
   c. Add 20 μL 200 mM NaIO4, mix and incubate 30 min at room temp. in the dark.
   d. Quench using 20 μL 50% ethylene glycol and incubate 30 min at room temp. in the dark.
   e. Buffer exchange using size exclusion spin columns.
      i. E.g. Micro Bio-Spin P-6 or Zeba 7K
      ii. First equilibrate column using 100 mM lysine
   f. Add 400 μL 1M lysine pH=8 + 1 μL SUPERase In.
   g. Incubate at 45°C for 5 hours.
   h. Purify using Monarch RNA Cleanup Kit (50 μg column version).
      i. RNA amount should be ~73 μg; therefore use two columns
      ii. The product is E.coli tRNA-Thr-CGT CCA-Phos
   i. Optional: gel purification of the product oligo.
   j. On a sample of the product oligo perform dephosphorylation to generate the E.coli tRNA-Thr-CGT CCA oligo.
   k. After RNA purification, measure RNA concentration and mix equal proportions of E.coli tRNA-Thr-CGT CCA-**Phos** and E.coli tRNA-Thr-CGT CCA to a final concentration of 20 ng/μL.
9. Optional: test buffers for RNase contamination using the RNaseAlert kit. This is optional since contamination will appear obvious during gel purification.

**A note about RNA stability**

Problems with RNA stability fall into two categories: RNase contamination and acid/base hydrolysis. We eliminate RNase contamination by general cleanliness such as using clean gloves, keeping surfaces and pipettes clean, using clean glassware and tube racks. We use RNaseZap (or equivalent) on suitable surfaces. We use Milli-Q purified water for making buffers over 10 mL in volume and find this to be sufficiently clean. For RNA elution/dilution we use nuclease free water. Occasionally, we use the RNaseAlert kit to test for contaminations. Finally, in some incubation steps an RNase inhibitor is used (as described below). In combination, we find that these steps make RNase contamination very unlikely.

Acid/base catalyzed RNA hydrolysis is inevitable but can be minimized. RNA stability is highest around pH 4-5, but it is the combination of pH and temperature that is important. Therefore, there is essentially only three steps in the RNA processing for the tRNA-charge sequencing protocol that can lead to RNA degradation: 1) the amine-induced cleavage at 45°C for 4 h, 2) the RNA denaturing at 90°C in loading buffer and 3) the gel elution at 65°C. Especially, for the amine-induced cleavage incubation step pH control is important. At pH=8 high RNA integrity is maintained while at pH=9.5 integrity is decreased substantially. The RNA denaturing step is performed at 90°C making the RNA extremely vulnerable to hydrolysis; therefore, we use a sodium acetate based loading buffer at pH=4.7 instead of the typical TBE based loading buffer at pH=8.3. Similarly, the gel elution is performed at low pH.

**Stepwise protocol**

**RNA oxidation, amine-induced cleavage and size selection**

1. Isolate total RNA using cold Trizol. Precipitate, wash twice with 80% IPA (containing 50 mM sodium acetate, pH=4.5), do a third wash with 100% IPA, then dry using centrivap or on the benchtop. Can be stored dry at -80°C for months without losing charge.
   a. Do not wash cells/tissue before adding Trizol as this will cause tRNA charge depletion.
   b. Everything must be processed within 2 hours and kept ice cold after adding Trizol to preserve the charge.
   c. Thorough washing is essential because Trizol contains glycerol, a quencher of NaIO4.
   d. Do not use a co-precipitant, as these often contain glycogen, a competing substrate for the subsequent NaIO4 oxidation step.
2. Reconstitute in 100 mM sodium acetate, pH=4.5 and keep on ice. Determine RNA concentration, adjust to 1 μg/μL and transfer 10 μL to a fresh tube. While keeping it cold, add 1 μL E.coli tRNA spike-in (optional) then add 5 μL 200 mM NaIO4 (prepared fresh), mix and incubate for 10 min on ice, in the dark. Then quench by adding 5 μL 50% (v/v) ethylene glycol (~9 M), mix and incubate 5 min on ice, in the dark. Place the tubes in a rack at room temperature for an additional 5 min before continuing.
   a. Should the RNA solution contain potassium ions, KIO4 will likely precipitate because of lower solubility. In such cases, increase the incubation time for oxidation to 30 min on ice, and for quenching 30 min on ice followed by 30 min at room temperature with mixing.
3. Perform amine-induced cleavage and deacetylation by adding 50 μL 1 M lysine (pH=8) with 1 μL SUPERase In (RNase inhibitor) and incubate for 4 h at 45°C.
4. Dephosphorylate RNA by adding 8 μL 10X rCutSmart Buffer and 1 μL rSAP and incubate for 30 min at 37°C.
5. Purify using Monarch RNA Cleanup Kit (50 μg column version), elute with 30 μL water, then perform size selection using polyacrylamide gel (see separate section).
   a. Use 6 μL eluted RNA of this for size selection.
   b. Cut out bands corresponding to tRNA (50 - 100 nt. range)
      i. Mature tRNAs vary in size from 62 to 90 nt.
      ii. Smallest rRNA is 5S rRNA at 118 nt.
      iii. Cut just below the middle between 80 nt. and 150 nt. (according to ladder) and/or just below the 5S rRNA band.
   c. Elute with 10 μL water and measure RNA concentration.
      i. Expect RNA concentration between 7-16 ng/μL

**Size selection using polyacrylamide gel**

1. Add RNA loading buffer to sample and heat denature 2 min at 90°C.
2. Run samples alongside NEB Low Range ssRNA ladder on a Novex 10% polyacrylamide/7M urea/1xTBE gel.
   a. Pre-run gel 10-40 min at 300V and thoroughly wash the wells with running buffer just prior to loading.
   b. Load, then run until the fastest tracking dye (bromophenol blue) reaches the bottom of the gel (~35 min at 300V).
      i. We run 10 min at 100V followed by 35 min at 300V.
      ii. Running the gel longer results in better resolution but also a larger gel volume to isolate RNA from, thus decreasing the yield.
   c. Incubate gel in TBE for 20 min to remove most of the tracking dye.
   d. Stain gel with SYBR Gold in TBE for 3 min, then transfer to fresh TBE.
   e. Visualize stain using a blue light transilluminator to avoid RNA cross linking.
   f. The tRNAs and rRNAs can often be observed as bands with a bright edge and dim center. This is due to fluorescent quenching because of the high density of RNA within the band.
3. Cut out the desired size range.
4. Add gel slice to a 1.5 mL tube and crush gel with disposable pestle. Then add 200 μL gel elution buffer + 1 μL Superase (for RNA samples only).
5. Snap freeze tubes with liquid nitrogen. Thaw and elute at 65°C for 5 mins with shaking.
6. Centrifuge samples through a SpinX centrifuge filter, purify using the Oligo purification kit (use 400 μL binding buffer and 1.1 mL EtOH for binding) and elute.

**Adapter ligation to tRNA 3’ ends**

1. Setup ligation reactions with:
   a. 4 μL gel-purified and annealed tRNA (35 ng total)
      i. Anneal in 1x NEBuffer 2 by heating to 94°C for 2 min followed by cooling 1°C/s to 4°C.
      ii. Avoid reheating annealed tRNA before ligation has finished.
   b. 1 μL 20 μM (20 pmol, ~10x excess) annealed adapter:splint partial duplex
   c. 2 μL 10x T4 RNA ligase buffer
   d. 4 μL 50% PEG-8000
   e. 1 μL NEBuffer 2
   f. 1 μL T4 RNA ligase 2
   g. 1 μL Superase In
   h. 6 μL water
2. Mix each reaction by pipetting and incubate 1 h at 37°C, then 24 h at 4°C
3. Heat inactivate 80°C for 5 min.
   a. Heat inactivate immediately prior to purification to avoid potential incubation in RNase released from the denatured RNase inhibitor.
4. Pool samples based on adapter barcodes and purify with the Oligo purification kit. Elute with 10 μL water.
5. Perform size selection using polyacrylamide gel (see separate section).
   a. Load the entire amount for size selection.
   b. Cut out range corresponding to adapter ligated tRNA (102 - 133 nt.)
      i. Make the lower cut at 90 nt. (according to the ladder). Including a small amount of unligated tRNA does not interfere with subsequent steps.
   c. Elute with 15 μL water and measure RNA concentration.

**RT PCR of adapter-tRNA**

1. In a PCR tube, combine:
   a. 10 μL adapter-tRNA (60 ng)
   b. 2 μL 1.25 μM primer-dependent RT oligo
   c. 4 μL RT buffer (5x First strand synthesis)
2. In addition to the samples, prepare an adapter only control with the shortest adapter. This serves as a size marker and can be prepared once and used several times.
3. Denature at 90°C for 2 min, anneal at 70°C for 30 seconds then cool 0.2°C/s to 4°C.
4. Then, to each tube add:
   a. 1 μL 100 mM DTT
   b. 1 μL Superase In
   c. 1 μL TGIRT-III
5. Mix and incubate at 42°C for 10 min.
6. Add 1 μL 25 mM dNTPs to each tube. Mix and incubate at 42°C for 16 h on a thermocycler with the heated lid set to 50°C.
7. Add 1 μL 5 M NaOH to each tube. Incubate at 95°C for 3 min to hydrolyze the RNA template.
8. Purify with the Oligo purification kit eluting with 10 μL water.
9. Perform size selection using polyacrylamide gel.
   a. Load the entire amount for size selection.
   b. Cut everything above the adapter only control.
      i. Full size tRNA is in the range 152-183 nt., no insert is 91-94 nt.
      ii. This intentionally includes truncated tRNA because of the incomplete readthrough of modified nucleotides.
   c. Elute with 7 μL water, no need to measure DNA concentration.

**cDNA circularization and library construction PCR**

1. Setup reactions in PCR tubes:
   a. 5.5 μL gel-purified cDNA
   b. 2 μL 5 M betaine
   c. 1 μL 10x CircLigase buffer
   d. 0.5 μL 1 mM ATP
   e. 0.5 μL 50 mM MnCl2
   f. 0.5 μL CircLigase
2. Mix each reaction by pipetting. Incubate at 60°C for 3 h on a thermocycler with a 70°C heated lid. Denature at 80°C for 10 min to inactivate the enzyme.
3. Use PCR to attach Illumina flanking sequences and barcodes. One PCR reaction contains:
   a. 0.6 μL circularized cDNA
   b. 1.5 μL 10 mM dNTPs
   c. 5 μL 10 μM library construction primer (P5)
   d. 5 μL 10 μM library construction primer (P7)
   e. 10 μL 5x KAPA HiFi buffer
   f. 1 μL KAPA HiFi Polymerase
   g. 26.9 μL water
4. Empirically, determine the appropriate amount of cycles by removing samples at three different cycle numbers. Amplify for with X cycles with the following settings:

| Temp. | Time | Cycles |
| --- | --- | --- |
| 95C | 3 min | 1 |
| 98C | 20 s | 3 |
| 68C | 10 s |  |
| 72C | 15 s |  |
| 98C | 20 s | X = 10, 12, 14 |
| 72C | 15 s |  |
| 4C | Inf | 1 |

5. After the PCR and onwards do not heat above room temperature to avoid reannealing. Reannealing will generate heteroduplexes which complicates subsequent gel isolation and quantification.
6. Find the best PCR settings by mixing 4 μL PCR product with 1 μL 5X non-denaturing sample loading buffer and run on a 4-12% non-denaturing TBE gel alongside the 10 bp DNA ladder. We run at 180V for 30 min.
7. For each sample, pool tubes with cycle numbers leading to specific amplification, then purify PCR products with the DNA cleanup kit, elute with 8 μL water and add 2 μL 5X non-denaturing sample loading buffer.
8. Run samples on a 4-12% non-denaturing TBE gel alongside the 10 bp DNA ladder.
   a. Run gel at 180V for 30-45 min.
   b. Stain with SYBR Gold in TBE for 3 min and isolate the 170-290 bp range (empty fragment is 165 bp) using a blue light transilluminator.
   c. Optional: Use adapter only (no insert) as a minimum size marker.
9. Add gel slice to a 1.5 mL tube and crush gel with disposable pestle. Then add 300 μL TBE.
10. Snap freeze tubes with liquid nitrogen. Thaw and elute at room temperature overnight with mixing.
11. Centrifuge samples through a SpinX centrifuge filter, purify DNA using the DNA cleanup kit. Elute with 20 μL 10 mM Tris pH=8 and quantify DNA concentration.
12. Optional: determine library length on TapeStation for some or all samples.
13. Use the estimated mean library length and the DNA concentration to calculate library molarity for each sample, then pool samples at equimolar concentration into the final sequencing library in a low binding tube.
14. Sequence using Illumina paired end sequencing using 2x100 bp reads.

**Gel at tRNA isolation step. Right, before, left, after size range cutout.**

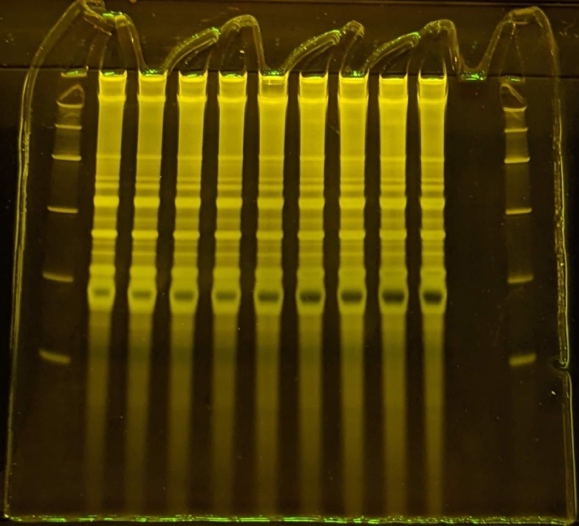

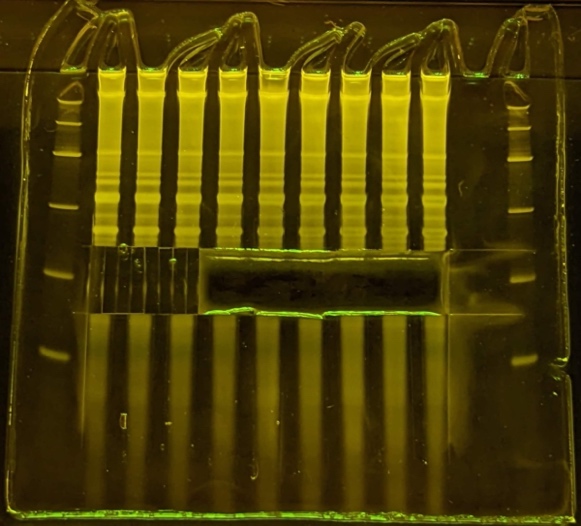

**Gel at adapter-tRNA isolation step. Right, before, left, after size range cutout.**

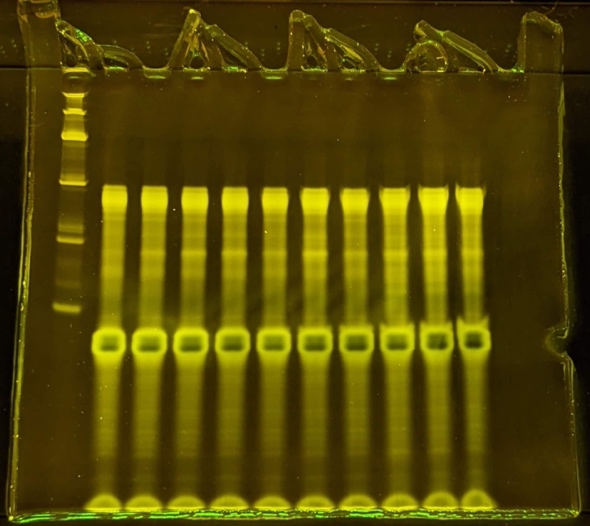

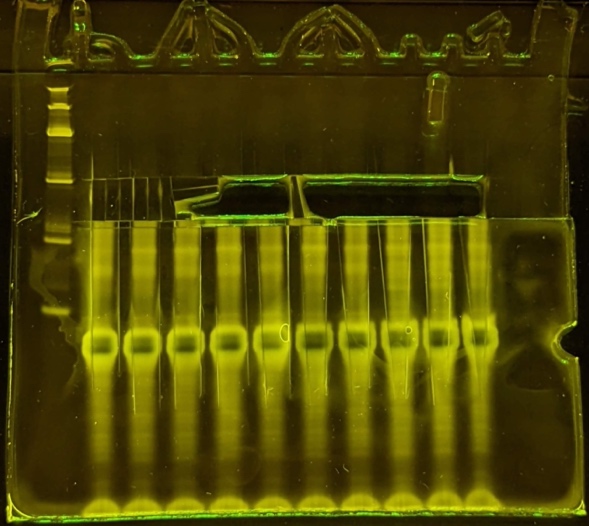

**Gel at cDNA isolation step. Right, before, left, after size range cutout.**

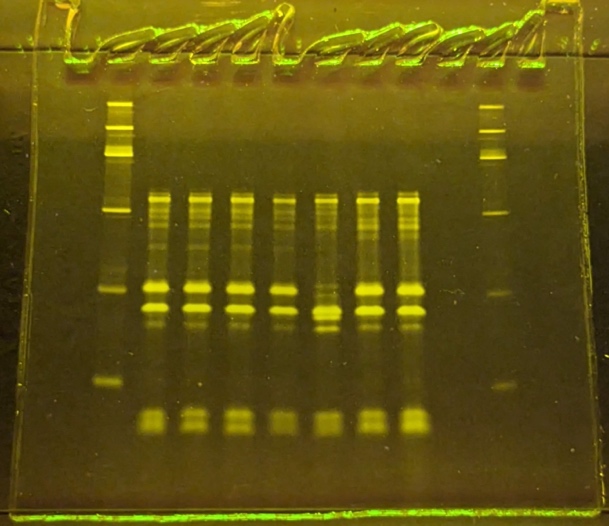

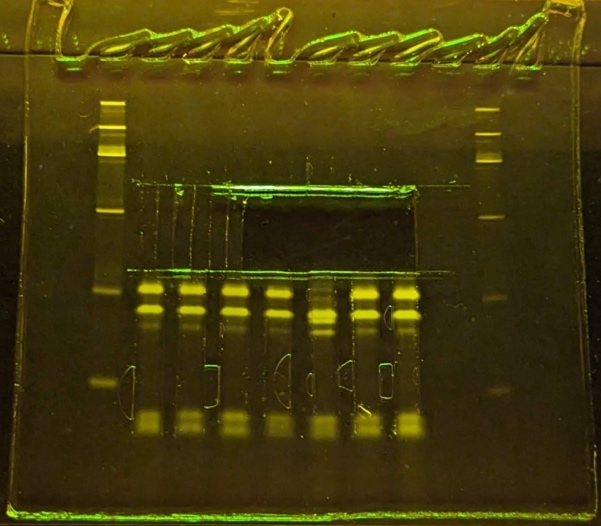

**Gel at library PCR step. From left to right, 10, 12 and 14 cycles, repeated for 4 samples.**

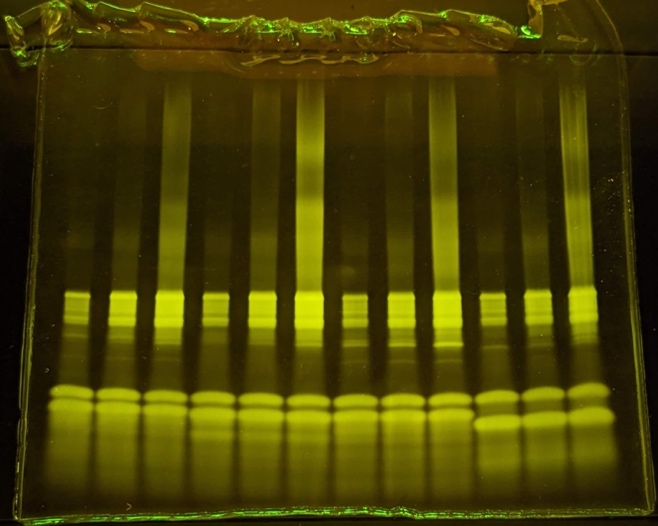

**Gel at library PCR step after pooling and purifying cycle 10 and 12 (see above). Right, before, left, after size range cutout.**

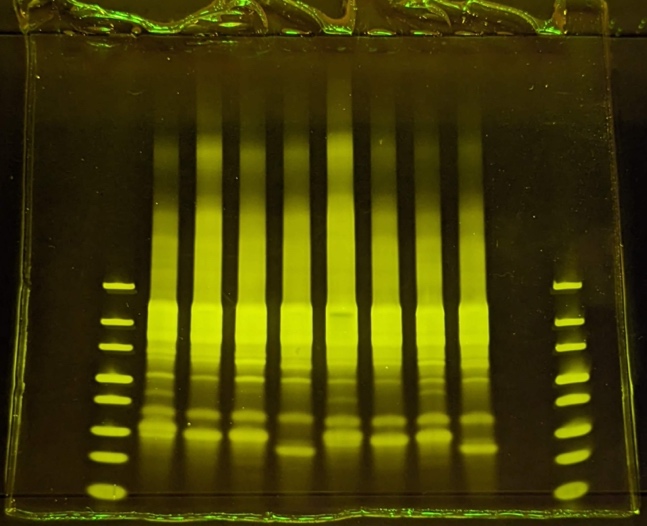

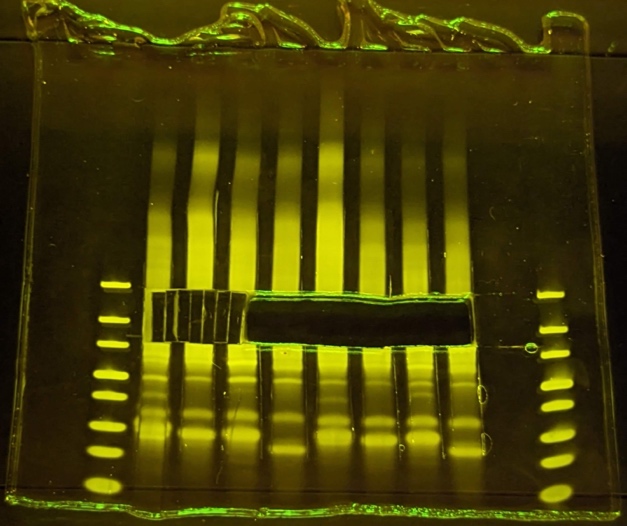
